# An Investigation of the Conformational Dynamics of ABC Exporter PCAT1 using Microsecond-Level MD Simulations

**DOI:** 10.64898/2026.03.04.709725

**Authors:** Matthew Brownd, Ehsaneh Khodadadi, Mahmoud Moradi

## Abstract

Peptidase-containing ATP-binding cassette transporters (PCATs) couple ATP hydrolysis with proteolytic processing and export of cargo peptides across cellular membranes. Despite their importance in bacterial secretion systems, the molecular determinants governing nucleotide binding and stabilization in PCAT transporters remain incompletely understood. In particular, recent experimental observations suggest that PCAT1 may display altered nucleotide preferences compared with canonical ABC transporters.

Here, we employed microsecond-scale all-atom molecular dynamics simulations combined with free energy perturbation (FEP) calculations to characterize nucleotide binding, protein stability, and conformational dynamics of PCAT1 across multiple biochemical conditions. Simulations were performed for inward-facing (IF) and outward-facing (OF) conformations in the presence or absence of Mg^2+^ and substrate peptides. Structural analyses reveal that substrate and Mg^2+^ jointly stabilize the IF conformation, reducing global structural fluctuations and enhancing nucleotide retention in the binding pockets. In contrast, systems lacking Mg^2+^ exhibit increased nucleotide mobility and partial dissociation events.

Thermodynamic analysis using FEP calculations further demonstrates that ATP binding is strongly stabilized in the IF state, particularly in the presence of Mg^2+^, whereas nucleotide stability is reduced when Mg^2+^ coordination is absent. To identify the molecular origins of nucleotide stabilization, we introduce a residue-level free energy decomposition approach that quantifies the contribution of individual residues to nucleotide binding energetics. This analysis reveals that the Walker A residue Lys525 provides the dominant stabilizing interaction with ATP, while neighboring residues within the Walker A motif contribute additional stabilization. In contrast, acidic residues of the Walker B motif primarily participate in catalytic organization rather than direct nucleotide stabilization.

Together, these results provide a comprehensive molecular description of nucleotide stabilization and conformational regulation in PCAT1. The combined structural and energetic analyses support a model in which Mg^2+^ coordination and substrate binding cooperatively stabilize the inward-facing state and organize the nucleotide-binding site for productive ATP hydrolysis. More broadly, this work demonstrates how residue-level free energy analysis can reveal the energetic architecture of nucleotide recognition in ABC transporters.

## Introduction

ATP-binding cassette (ABC) transporters are integral membrane proteins that couple the free energy of ATP binding and hydrolysis to the directional transport of chemically diverse substrates across cellular membranes.^1–7^ ABC transporters are found in all domains of life and participate in processes ranging from nutrient uptake and lipid trafficking to drug efflux and peptide secretion.^3,5^ Despite this functional diversity, the core architecture is conserved: two transmembrane domains (TMDs) form the substrate translocation pathway, and two nucleotide-binding domains (NBDs) bind/hydrolyze ATP to drive large-scale conformational rearrangements in the TMDs.^2,4^

ABC exporters generally operate via an alternating-access mechanism in which the central cavity alternates between inward-facing (IF) and outward-facing (OF) conformations.^5,6^ In the canonical picture, ATP binding promotes NBD dimerization and stabilizes an OF state that enables substrate release; ATP hydrolysis and product release then reset the transporter toward an IF conformation that is competent for another round of substrate binding.^5,7^ Structural studies of bacterial exporters such as Sav1866 and MsbA were instrumental in establishing this coupling paradigm, showing how nucleotide-state-dependent changes at the NBD interface reorganize the TMD helices and reshape the substrate cavity. ^8,9^

Among ABC exporters, the protease-containing ABC transporters (PCATs) represent a specialized subfamily responsible for processing and exporting polypeptide cargos such as bacteriocins and quorum-sensing peptides, central to bacterial defense and communication mechanisms.^10–13^ Many bacteriocins are ribosomally synthesized as precursor peptides bearing N-terminal leader sequences that prevent premature activity and guide maturation and secretion.^14^ A defining feature of PCAT systems is an N-terminal C39 peptidase domain fused to the ABC core, which cleaves the leader peptide prior to or during translocation, thereby coupling proteolytic processing and membrane export within a single molecular machine.^11,15^

PCAT1, a well-characterized member from *Clostridium thermocellum*, has emerged as a key model for understanding how PCAT transporters coordinate peptide recognition, processing, and extrusion.^11,13^ Recent cryo-EM studies have captured PCAT1 in multiple conformational states along its transport cycle, including inward-facing (IF), occluded (OC), and outward-facing (OF) conformations.^11,16,17^ These structures provided an initial mechanistic framework for how peptide substrates engage the transporter and how the C39 peptidase domain is positioned relative to the transmembrane channel during maturation and export.

Nevertheless, the mechanistic coupling between nucleotide chemistry at the NBDs and cargo processing/transport in PCATs is not fully understood.^18^ Compared with canonical ABC exporters, PCATs must coordinate additional regulatory steps, including proteolytic processing of a polypeptide substrate, suggesting a more intricate interplay between ligand binding, conformational dynamics, and substrate recognition.^19^ In this context, nucleotide-state energetics are particularly important because ATP, ADP, and Mg^2+^ binding can reshape the free-energy landscape of conformational interconversion, influencing which conformations are populated under physiological conditions.

Recent cryo-EM and biochemical analyses suggest that PCAT1 exhibits unusually high affinity for ADP compared to ATP, particularly in the OF state, which is contrary to the binding trends observed in most bacterial ABC transporters.^19^ This reversed preference has been proposed to serve a regulatory function, preventing futile ATP hydrolysis in the absence of substrate. Moreover, structural and functional data implicate the substrate as a modulator of nucleotide binding, capable of reversing ADP-mediated inhibition and stabilizing the IF conformation.^20,21^ Together, these observations point to strong thermodynamic coupling between substrate binding, divalent cation coordination, nucleotide identity, and the global conformational state of PCAT1.

To quantitatively assess these observations, we performed all-atom molecular dynamics (MD) simulations and free energy perturbation (FEP) calculations to determine the relative binding affinities of ATP and ADP in PCAT1 across multiple conformational states (IF, OC, OF) and biochemical conditions (±Mg^2+^, ±substrate). Our simulations also provide per-residue energetic insights into key structural motifs, such as the A-loop and signature motif, implicated in nucleotide binding and allosteric coupling. By integrating free-energy calculations with available structural data, this work provides a quantitative framework for understanding how PCAT1 selectively couples ATP hydrolysis to peptide processing and export.

## Methods

All-atom molecular dynamics (MD) simulations were performed on multiple systems of the peptidase-containing ABC transporter PCAT and the bacterial transporter MsbA. An ATP-bound, outward-facing cryo-EM structure of PCAT (PDB: 7T54) served as the template for system construction. Magnesium ions were introduced by aligning the ATP ligands of this structure with those from PDB: 7T57, which contains Mg^2+^-coordinated nucleotides. The aligned nucleotides from 7T57 were substituted into the PCAT structure to incorporate magnesium ions appropriately.

Three conformational states of MsbA (inward-facing closed, inward-facing occluded, and outward-facing) were similarly prepared as control systems. These models, derived from prior work by Moradi and Tajkhorshid, were ATP- and Mg^2+^-bound by alignment with nucleotide-binding domains of known structures. Parallel systems lacking Mg^2+^ were generated by selectively removing magnesium ions from the models. This resulted in eight systems (four conformational states each with and without Mg^2+^), enabling comparative binding free energy analysis. Two additional systems were prepared containing only ATP, with or without Mg^2+^, solvated in water, serving as control systems for the free-energy perturbation (FEP) calculations.

A second set of PCAT systems, both with and without Mg^2+^, was constructed using the same 7T54 template. Missing loop residues (151–157) were modeled into both protomers using Modeller^22,23^ to complete the structure. These systems were used both for residue-specific interaction free energy calculations and extended equilibrium MD simulations on Anton.

The third and fourth sets of systems consisted of nucleotide- and substrate-bound PCAT models provided by collaborators. Five nucleotide-bound PCAT structures (e.g., PCAT1 CL IF ADP noSu PCAT1 OC ATP noSub) were retained with their original nucleotide and Mg^2+^ states, while substrates were removed for simplification. These systems were simulated under equilibrium conditions. Additionally, six substrate-bound PCAT systems were constructed (e.g., PCAT1 IF ATP Sub Mg, PCAT1 IF ADP Sub), and ATP or Mg^2+^ was introduced or removed as necessary through alignment with PDB: 7T54.

All systems were processed through the CHARMM-GUI Solution and Membrane Builders,^24–26^ using PPM2.0 to orient the transporter within a lipid bilayer composed of 45% POPE, 45% POPG, and 10% cardiolipin (TOCL2). Systems were solvated using TIP3P water^27^ with 0.15 M NaCl and appropriate counterions to ensure electroneutrality. Each complete simulation system contained approximately 450,000 atoms.

Simulations were carried out using NAMD 2.14^28,29^ with the CHARMM36m force field for proteins and ions. ^30^ The ATP and ADP ligands were parameterized using standard CHARMM protocols. Each system underwent energy minimization via the conjugate gradient algorithm for 10,000 steps, followed by a six-step restraining protocol over 18.75 ns in the NVT ensemble. Production runs were performed for 20 ns under NPT conditions at 310 K using Langevin dynamics.^31^ Pressure was maintained at 1 atm using the Nosé–Hoover Langevin piston method.^32,33^ Long-range electrostatics were handled with the particle mesh Ewald (PME) method,^34,35^ and nonbonded interactions were truncated at 12 ^å^.

Extended equilibrium simulations for selected PCAT systems were performed on the Anton supercomputer,^36,37^ using structures equilibrated as described above. Each trajectory was run for 1.2 *µ*s with a 2.5 fs time step. Temperature was maintained at 310 K using the Nosé–Hoover thermostat,^38,39^ and pressure was controlled semi-isotropically at 1 atm using the MTK barostat.^33^ Long-range electrostatics were computed via the FFT-based PME method implemented on Anton.

Free-energy perturbation (FEP) simulations were conducted to estimate the relative binding free energy of ATP/ADP and to determine per-residue contributions. The FEP process modeled the hydrolysis of ATP to ADP by alchemically annihilating the *γ*-phosphate group of the nucleotide, following the standard free-energy perturbation formalism.^40^ This was done in both Mg^2+^-bound and unbound conditions across 20 *λ*-windows (Δ*λ* = 0.05). Each window consisted of 10 ps of equilibration followed by 240 ps of production sampling, for a total of 5 ns per FEP trajectory. All windows were initialized from equilibrated structures with partially annihilated phosphates, allowing simultaneous parallel execution. FEP simulations were performed using NAMD 2.13 ^28^ with the CHARMM36m force field^30^ and a 1 fs timestep. Soft-core potentials were applied to address endpoint issues in van der Waals decoupling, with electrostatic interactions fully removed by *λ* = 0.5. Free energy differences were calculated using the Bennett Acceptance Ratio (BAR) method.^41^ VMD’s ParseFEP plugin was used to analyze FEP results.^42^

Interaction free energies between individual residues and ATP/ADP were calculated from the FEP trajectories using NAMD’s pairinteraction module. Residues from conserved functional motifs were examined, including the Walker A motif (G519–T526 in PCAT), the Walker B motif (I643–E648), and the ABC signature motif (L623–Q627). Electrostatic and van der Waals energy contributions were sampled every 5 ps (1000 points per FEP trajectory). For each window, the equilibrium constant *K*_window*,i*_ was calculated using Boltzmann-weighted averages of energy differences. Different equations were applied for electrostatics-dominant (windows 1–10) and van der Waals-dominant (windows 11–20) phases. The total interaction free energy (Δ*G*) for each residue was then computed as:

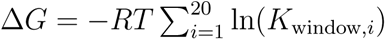

These values were used to evaluate the significance of individual residue contributions to ATP/ADP binding under varying magnesium conditions.

## Results and Discussion

### Ligand RMSD Reveals Conformational Stability Across PCAT1 Systems

To assess the structural stability of ligand binding across different PCAT1 systems, we analyzed the root-mean-square deviation (RMSD) of the ligand relative to its initial bound pose throughout the simulation trajectories. Ligand RMSD provides a direct measure of how much the nucleotide deviates from its starting configuration and therefore offers insight into the stability of the binding mode over time (i.e., whether the bound pose remains well retained or instead explores alternative local conformations).

As shown in Figure 1, two representative systems demonstrate distinct patterns of ligand stabilization. In both cases, RMSD increases during the early phase of the simulation, reflecting initial relaxation from the modeled pose as the nucleotide adapts to the local geometry of the binding pocket. One system (left panel) quickly reaches a stable plateau with low-amplitude fluctuations, consistent with a tightly retained ligand and a binding pocket that rapidly equilibrates around it. The other system (right panel) exhibits a slower rise and sustained higher RMSD values, suggesting either slower convergence or enhanced flexibility of the ligand and/or binding pocket, allowing sampling of multiple nearby configurations.

**Fig. 1.**
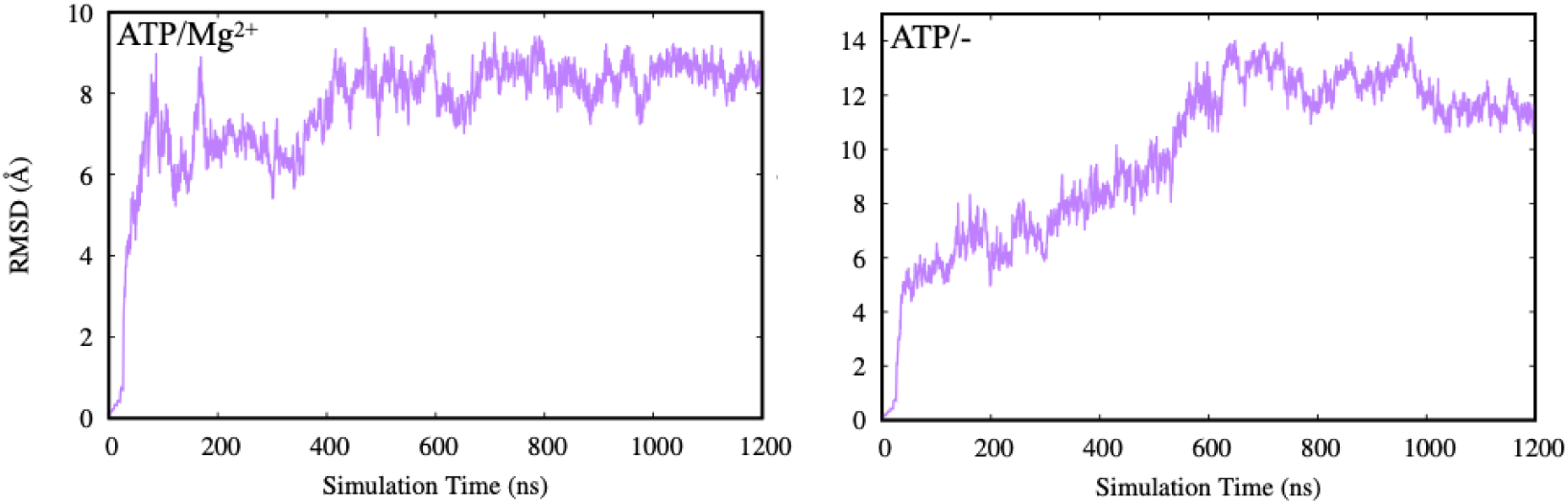
Representative ligand RMSD trajectories from two PCAT1 systems. In both cases, the ligand undergoes an initial relaxation period, followed by stabilization. The left panel shows rapid convergence to a low RMSD plateau, indicating strong retention of the ligand in the binding site. The right panel exhibits higher RMSD values and delayed stabilization, suggesting increased ligand mobility or slower equilibration

Importantly, elevated RMSD does not necessarily imply complete unbinding; rather, it can reflect local rearrangements such as changes in phosphate orientation, altered coordination of Mg^2+^ (if present), or small shifts in loop and motif positioning around the nucleotide. These modes of variability are particularly relevant for ABC transporters, where nucleotide-binding-site geometry is coupled to global conformational changes of the NBDs and the TMDs.

Figure 2 displays RMSD traces from four simulation replicates across distinct biochemical conditions. The top two replicates maintain low and steady RMSD values around 2–3 ^å^, indicating stable ligand binding. In contrast, the bottom two replicates display increased fluctuations or progressive drift, which may signal a degree of binding-site rearrangement, transient weakening of key contacts, or sampling of alternate nucleotide poses. Such differences can arise from (i) distinct starting conformations (IF versus OC-like), (ii) the presence/absence of substrate and Mg^2+^, and (iii) the degree of NBD closure and inter-domain coupling during the trajectory.

**Fig. 2.**
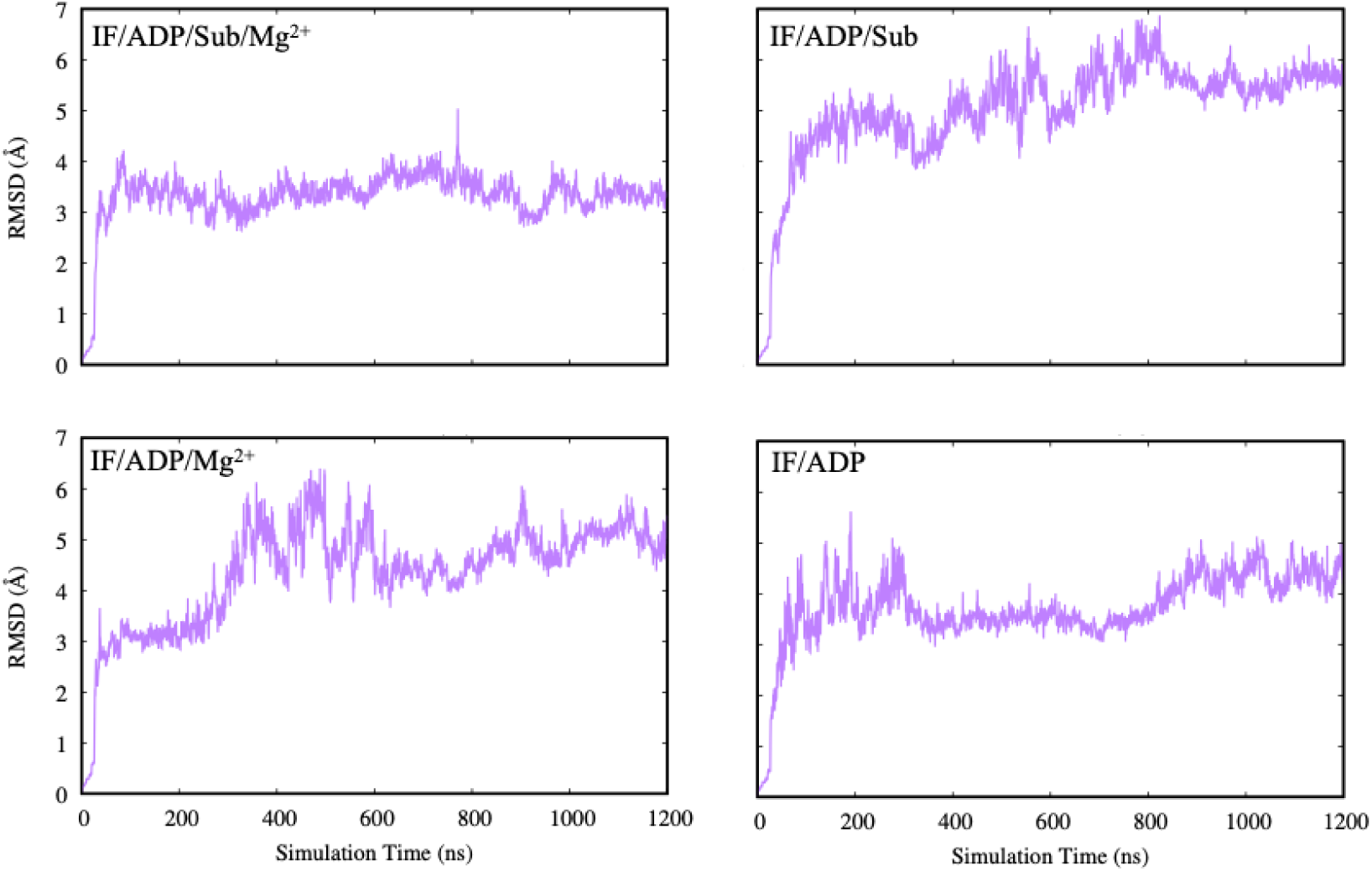
Root mean square deviation (RMSD) of C*α* atoms for PCAT1 in the inward-facing (IF) conformation under four ligand conditions: IF/ADP (bottom right), IF/ADP·Mg^2+^ (bottom left), IF/ADP/Sub (top right), and IF/ADP/Sub·Mg^2+^ (top left). The IF/ADP/Sub·Mg^2+^ system shows the lowest fluctuation and highest structural stability, while systems lacking either substrate or Mg^2+^ exhibit elevated RMSD values and greater dynamic variability.

Recent structural work has demonstrated that the conformational landscape of PCAT1 is highly responsive to the identity and occupancy of bound nucleotides, magnesium ions, and substrate peptides. ADP in the presence of substrate favors inward-facing conformations that expose a cytoplasmic chamber and stabilize substrate engagement. By contrast, ATP binding—particularly when Mg^2+^ coordination is compromised—can shift the local nucleotide geometry and alter the stability of the binding pocket, potentially biasing the system toward occluded or transitional states. Consistent with this model, the trajectories with low ligand RMSD are more compatible with a well-organized IF-like nucleotide-binding site, whereas the higher-RMSD traces likely reflect increased dynamical rearrangements within the binding pocket and associated structural elements.

Overall, the ligand RMSD analysis provides an initial, trajectory-level view of how biochemical conditions and conformational context influence nucleotide stability in PCAT1. In the following sections, we connect these qualitative stability patterns to global protein stability (RMSD), local flexibility (RMSF), direct measures of nucleotide retention, and ultimately thermodynamic estimates from free-energy calculations.

### Structural Stability of PCAT1 in the IF Conformation

To assess the overall structural stability of PCAT1 in the inward-facing (IF) conformation, we computed the root mean square deviation (RMSD) of the C*α* atoms across four simulation conditions: IF/ADP, IF/ADP·Mg^2+^, IF/ADP/Sub, and IF/ADP/Sub·Mg^2+^. Backbone RMSD reports on global conformational drift and provides a useful complement to ligand RMSD, allowing us to distinguish ligand instability arising from local binding-site rearrangements versus broader protein-level conformational changes.

As shown in Figure 2, all systems undergo an initial relaxation phase within the first ∼100 ns, followed by relatively stable RMSD profiles. This early increase likely reflects equilibration of the membrane-protein-solvent environment and modest readjustments of the NBD–TMD interface as the system relaxes from its initial modeled configuration. After this equilibration phase, each trajectory fluctuates around a characteristic RMSD range, indicating that the transporter remains globally stable on the simulated timescale.

However, distinct differences in fluctuation patterns emerge depending on ligand coordination. The IF/ADP/Sub·Mg^2+^ system (top left) exhibits the most stable behavior, with RMSD values remaining close to 3 ^å^ throughout the trajectory. This suggests that the combination of substrate and Mg^2+^ produces a particularly well-stabilized IF ensemble, consistent with a cooperative stabilization mechanism in which substrate engagement helps organize the TMD and coupling helices while Mg^2+^ reinforces catalytic-site geometry and NBD organization.

In contrast, the IF/ADP/Sub and IF/ADP·Mg^2+^ simulations (top right and bottom left) show higher variability and gradual structural deviation, particularly between 400–800 ns. One interpretation is that removing either Mg^2+^ or substrate weakens one stabilizing axis of the IF state, allowing larger amplitude breathing motions between NBDs and/or at the TMD–NBD coupling interface. The IF/ADP condition (bottom right) stabilizes after ∼400 ns but maintains slightly higher RMSD fluctuations compared to the fully coordinated system, again consistent with an IF ensemble that is stable but less tightly constrained.

Together, these results indicate that substrate binding and Mg^2+^ coordination act synergistically to maintain conformational integrity of PCAT1 in the IF state. This is mechanistically reasonable: Mg^2+^ directly stabilizes phosphate coordination and catalytic-site architecture, while substrate binding can allosterically stabilize coupling elements that connect the NBDs and TMDs, thereby reducing global conformational wandering that could otherwise destabilize the nucleotide-binding environment.

### Structural Flexibility of PCAT1 in the IF Conformation under ADP and Mg^2+^ Conditions

To evaluate how nucleotide, substrate, and Mg^2+^ binding affect local dynamics, we analyzed the root mean square fluctuations (RMSF) of C*α* atoms across the same four IF conditions (Figure 3). RMSF provides a residue-level map of flexibility and is therefore useful for identifying loop regions or peripheral segments that respond to ligand coordination, even when the global fold remains stable (as indicated by RMSD).

**Fig. 3.**
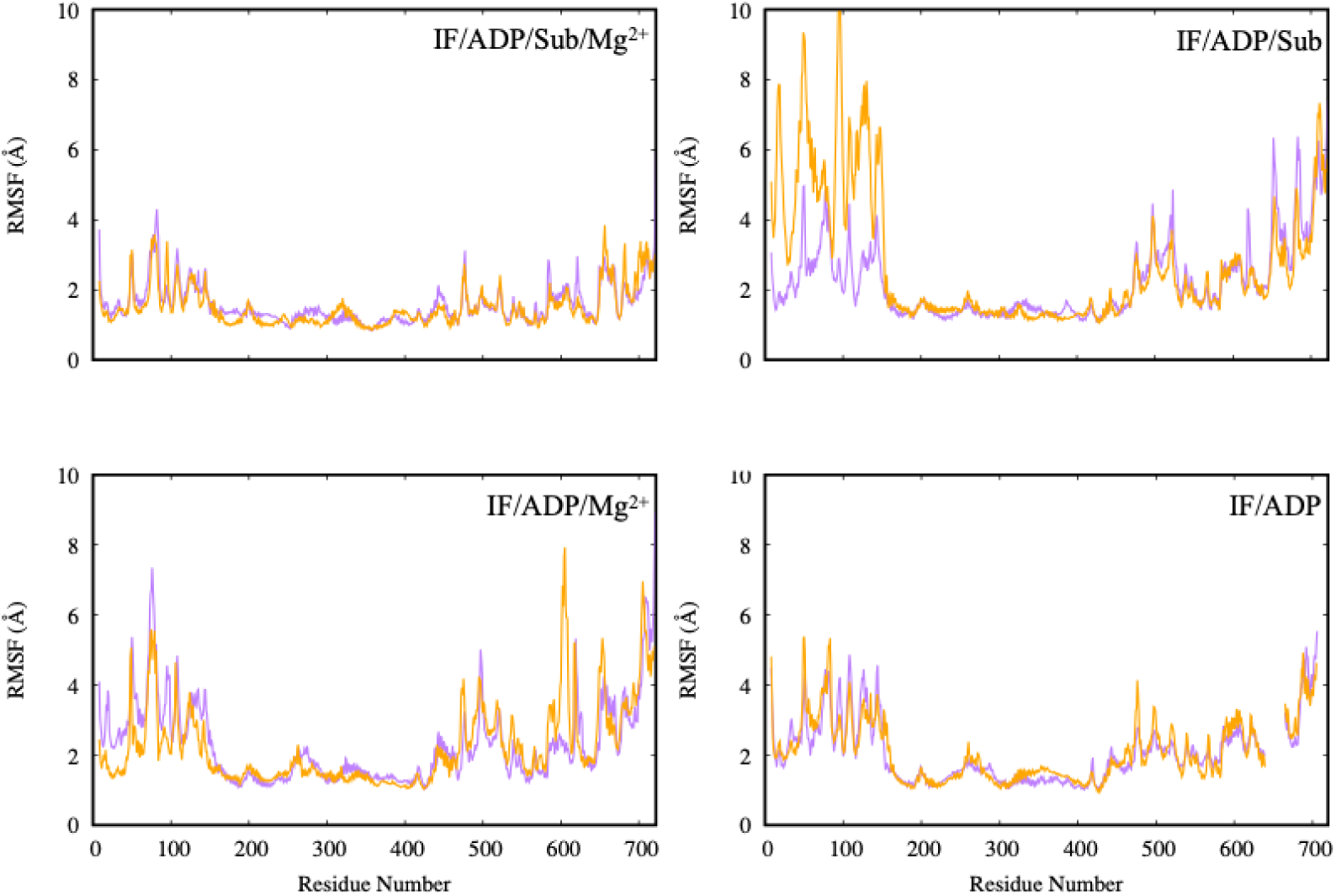
Root mean square fluctuations (RMSF) of backbone C*α* atoms across four inward-facing (IF) PCAT1 simulations: IF/ADP (bottom right), IF/ADP·Mg^2+^ (bottom left), IF/ADP/Sub (top right), and IF/ADP/Sub·Mg^2+^ (top left). The highest flexibility is observed in the IF/ADP/Sub system, particularly in loop and N-terminal regions. Addition of Mg^2+^ reduces these fluctuations, suggesting a stabilizing role for divalent cations in substrate-bound and nucleotide-bound states.

As shown in Figure 3, the IF/ADP/Sub simulation (top right) displays the highest degree of fluctuation in the N-terminal region and several loop segments, suggesting increased flexibility in the absence of Mg^2+^. This trend is mitigated in the presence of Mg^2+^: the IF/ADP/Sub·Mg^2+^ simulation (top left) shows a marked reduction in RMSF values across these regions, indicating enhanced rigidity when both substrate and Mg^2+^ are present.

Similarly, the IF/ADP simulation (bottom right) shows elevated flexibility compared to IF/ADP·Mg^2+^ (bottom left), reinforcing the stabilizing role of divalent cation coordination. Across all conditions, fluctuations cluster primarily in loop and peripheral regions, while the transmembrane helices and core remain relatively stable. This partitioning is consistent with a model in which the transporter’s structural scaffold remains intact while flexible elements modulate access, coupling, and potentially the kinetics of transitions between IF and downstream states.

Notably, enhanced flexibility in the absence of Mg^2+^ is consistent with the idea that Mg^2+^ does more than simply stabilize phosphate coordination; it can also indirectly reduce dynamical disorder in surrounding motifs and loops by reinforcing the local electrostatic network within the NBDs. These observations align with recent experimental evidence^19^ suggesting that nucleotide and substrate interactions regulate local flexibility, which may in turn influence the timing of ATP hydrolysis and conformational transitions in ABC exporters.

### Ligand Stability in the IF Conformation Under ADP and Mg^2+^ Conditions

While ligand RMSD reports on pose stability, it does not directly report on whether the nucleotide remains bound. To more directly assess nucleotide retention, we calculated the center-of-mass distance between each ADP molecule and its respective binding pocket across the four IF conditions (Figure 4). In this analysis, sustained low distances indicate stable binding, whereas large increases indicate partial or complete dissociation.

**Fig. 4.**
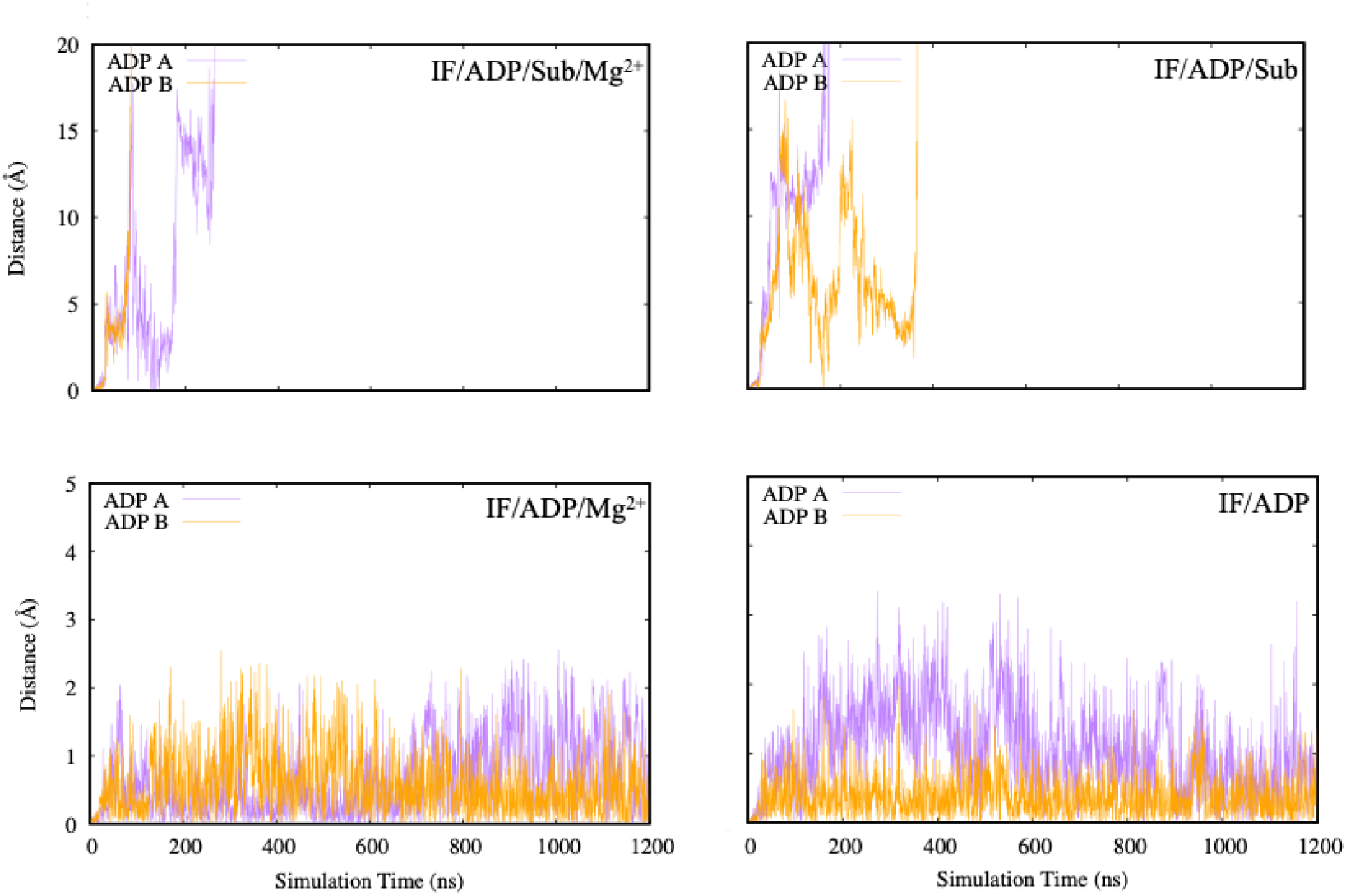
Center-of-mass distances between ADP and the nucleotide-binding pocket in four inward-facing (IF) PCAT1 simulation conditions: IF/ADP (top left), IF/ADP·Mg^2+^ (top right), IF/ADP/Sub (bottom left), and IF/ADP/Sub·Mg^2+^ (bottom right). In the absence of magnesium, ADP becomes unstable and often dissociates from one or both binding sites. The presence of Mg^2+^, and particularly the combination of Mg^2+^ and substrate, leads to consistent nucleotide retention, indicating a stabilizing effect of ligand coordination.

As shown in Figure 4, the presence of magnesium ions is critical for stabilizing ADP within the nucleotide-binding sites. In both IF/ADP and IF/ADP/Sub simulations (top left and bottom left panels), one or both ADP molecules progressively dissociate, with center-of-mass distances reaching values above 10 ^å^. This indicates weak or transient binding in the absence of divalent cations, consistent with the role of Mg^2+^ in coordinating phosphate groups and supporting the catalytic-site electrostatic network. These trajectories suggest that, without Mg^2+^, ADP may remain only loosely associated and can be displaced by local fluctuations in the binding pocket.

Conversely, simulations that include Mg^2+^ (top right and bottom right panels) show stable retention of ADP throughout the trajectory, with distances consistently below ∼2 ^å^. The addition of substrate in the IF/ADP/Sub·Mg^2+^ system further reinforces ADP stability, with minimal fluctuations in distance. This finding is consistent with experimental evidence that substrate interaction with regions near the A-loop can allosterically influence nucleotide stabilization by altering the local structural environment of the nucleotide-binding sites.^19^

Taken together, these results support a model in which ADP exhibits high-affinity binding only when Mg^2+^ is present, and that substrate binding can further enhance nucleotide retention by stabilizing the IF architecture. In functional terms, such an ADP-stabilized IF state could act as a regulatory checkpoint that prevents premature cycling and ensures that energy expenditure is tightly coupled to productive substrate processing and export.

### Free Energy Perturbation Reveals Ligand Stability Across Conformations

To quantify nucleotide binding thermodynamics across conformational and biochemical states of PCAT1, we employed free energy perturbation (FEP) calculations to compute ATP binding free energies under multiple conditions (Table 1). This analysis enables comparison of ATP affinity across IF versus OF conformations as well as across Mg^2+^ and substrate states, providing a thermodynamic complement to the structural stability metrics above.

**Table 1:**
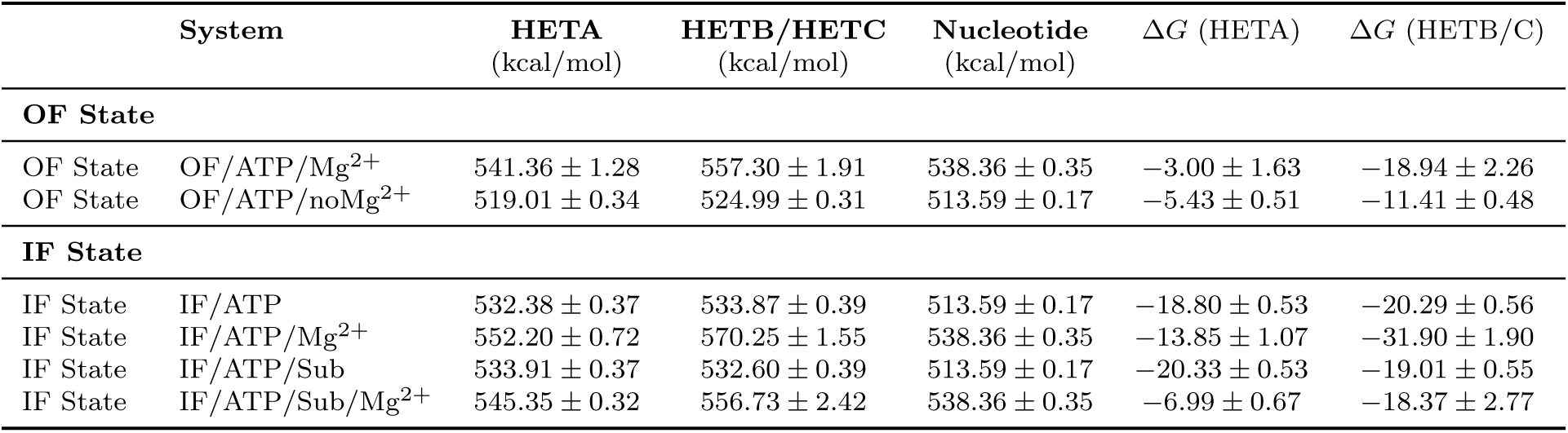
Free energy perturbation (FEP) calculations reveal ATP binding affinities under various PCAT1 simulation conditions. Shown are the energy values for the HETA and HETB/HETC sites, nucleotide reference energies, and corresponding relative binding free energies (Δ*G*). The data illustrate that ATP binding is highly stabilized in the inward-facing state, particularly when Mg^2+^ and substrate are both present. In the outward-facing conformation, Mg^2+^ also contributes significantly to nucleotide retention, but less so than in the inward-facing states.

In the outward-facing (OF) conformation, ATP binding is moderately stabilized in the presence of Mg^2+^, with a Δ*G* of −18.94 ± 2.26 kcal/mol at the HETB/C site. This stabilization is significantly reduced when Mg^2+^ is absent (Δ*G* = −11.41 ± 0.48 kcal/mol), underscoring the essential role of divalent cations in enhancing nucleotide affinity in the OF state. The site dependence (HETA versus HETB/C) suggests that local pocket geometry and inter-domain contacts differ between the composite sites, and that these differences can be amplified by Mg^2+^ coordination.

In the inward-facing (IF) state, ATP exhibits stronger binding and a more pronounced dependence on biochemical context. ATP binding in IF/ATP is favorable at both sites (Δ*G* = −18.80±0.53 kcal/mol at HETA and −20.29±0.56 kcal/mol at HETB/C), consistent with a binding environment that is compatible with stable nucleotide coordination. Upon addition of Mg^2+^, the HETB/C site becomes substantially more stabilized (Δ*G* = −31.90±1.90 kcal/mol), highlighting the strong energetic contribution of Mg^2+^ to phosphate coordination and to organizing catalytic-site residues. Interestingly, the corresponding HETA value becomes less favorable (−13.85 ± 1.07 kcal/mol), which may reflect asymmetry between the composite sites and/or conformational heterogeneity that differentially affects the two pockets.

Substrate binding also impacts ATP affinity. In IF/ATP/Sub, the HETA site is slightly more favorable than IF/ATP (−20.33 ± 0.53 kcal/mol versus −18.80 ± 0.53 kcal/mol), suggesting substrate-induced stabilization of at least one binding pocket. However, the triply coordinated IF/ATP/Sub/Mg^2+^ system does not yield the most favorable Δ*G* values, which may reflect entropic penalties, steric constraints, or substrate-dependent shifts in the conformational ensemble that reshape the binding environment in a non-additive manner. In other words, Mg^2+^ and substrate do not necessarily contribute independently; instead, the combined state may induce structural adjustments that partially offset raw electrostatic stabilization.

Overall, the FEP results support a model in which ATP affinity depends strongly on both transporter conformation and biochemical context, consistent with ligand-induced conformational selection. Importantly, these thermodynamic trends provide a quantitative basis for interpreting how PCAT1 may regulate ATP binding/hydrolysis as a function of state and substrate occupancy.

### Residue-Level Free Energy Decomposition Reveals Determinants of Nucleotide Recognition

To identify the molecular origins of nucleotide stabilization in PCAT1, we performed a per-residue decomposition of the interaction free energy between ATP and residues in the nucleotide-binding pocket (Table 2). This analysis represents a novel methodological component of the present study, in which the contribution of individual residues to the overall nucleotide-binding free energy is quantified directly from the free energy perturbation (FEP) trajectories. In this framework, the reported values reflect the relative energetic contribution of each residue to the stabilization of the *γ*-phosphate group of ATP during the ATP→ADP alchemical transformation.

**Table 2:**
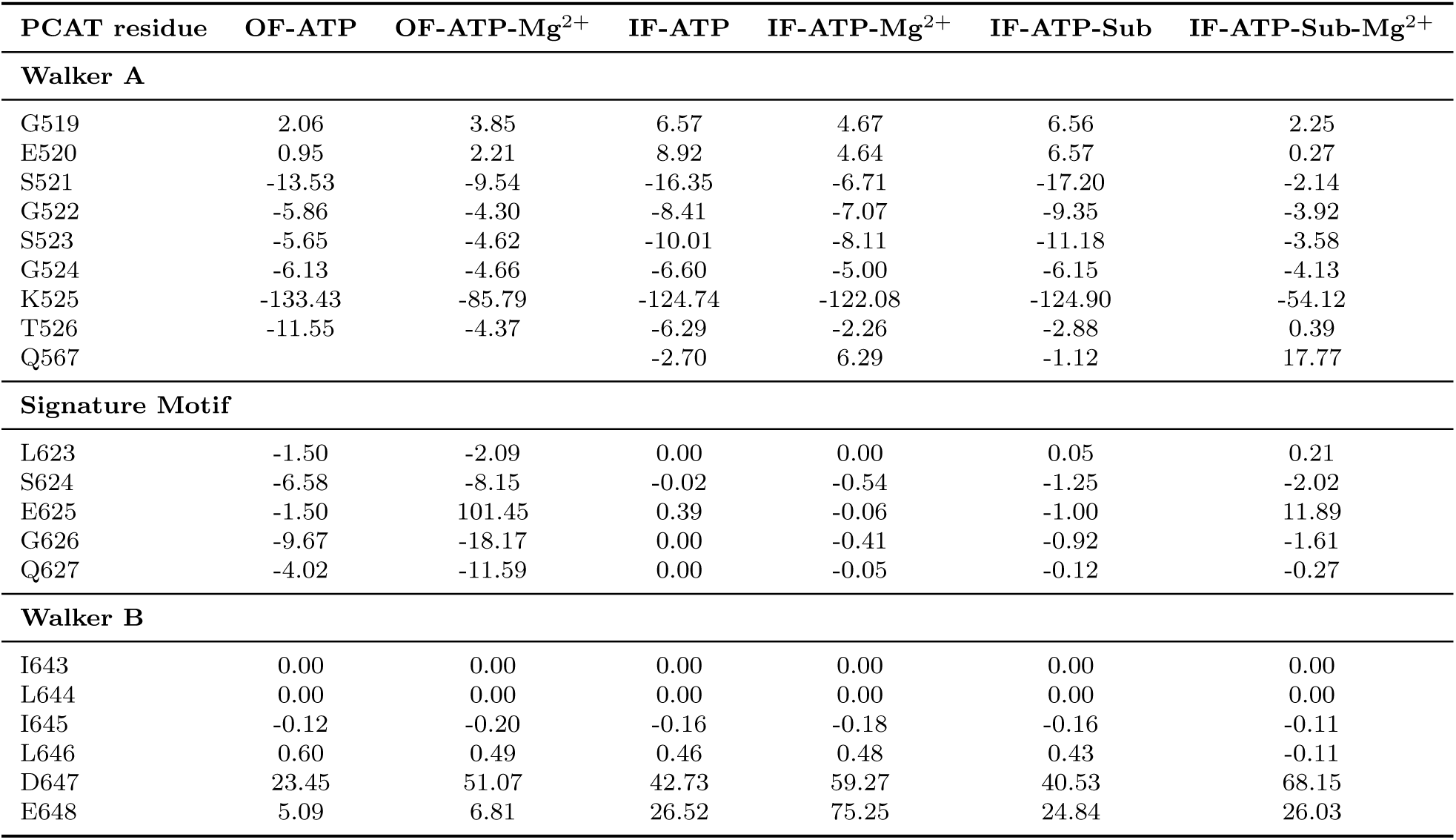
Relative per-residue interaction free energy with the γ-phosphate (kcal/mol).

The resulting energy decomposition reveals a clear hierarchy of residue contributions within the conserved nucleotide-binding motifs of PCAT1. As expected, the largest stabilizing interaction arises from Lys525 within the Walker A motif. Across all simulated conditions, Lys525 exhibits the strongest negative interaction energy, reaching values as large as −133.43 kcal/mol in the OF-ATP system and −124.74 kcal/mol in the IF-ATP system.

This strong contribution reflects the well-established role of the Walker A lysine in coordinating the phosphate groups of ATP through electrostatic interactions. The magnitude of this interaction highlights the central role of Lys525 in anchoring the nucleotide within the binding pocket.

Additional stabilizing contributions are observed for several neighboring residues within the Walker A motif, including Ser521, Gly522, Ser523, and Gly524. These residues consistently display negative interaction energies, indicating favorable interactions with the nucleotide phosphates. In particular, Ser521 and Ser523 show strong stabilizing contributions across multiple simulation conditions, suggesting that hydrogen-bonding interactions involving the backbone and side-chain hydroxyl groups play an important role in stabilizing ATP within the pocket.

Residues in the ABC signature motif contribute more modestly to nucleotide stabilization. Among these residues, Ser624 and Gly626 display consistently favorable interaction energies, although the magnitudes are smaller than those observed for Walker A residues. This pattern is consistent with the structural role of the signature motif in mediating interdomain contacts between the two nucleotide-binding domains rather than directly coordinating the nucleotide.

In contrast, residues in the Walker B motif exhibit positive interaction energies in several conditions, most notably Asp647 and Glu648. These residues show large positive values, reaching up to 59.27 kcal/mol for Asp647 and 75.25 kcal/mol for Glu648 in the IF-ATP-Mg^2+^ condition. Positive interaction energies indicate that these residues do not directly stabilize the nucleotide; instead, they likely participate in catalytic positioning and coordination of Mg^2+^ during ATP hydrolysis. In canonical ABC transporters, the Walker B acidic residues play a key role in activating the catalytic water molecule rather than directly stabilizing the nucleotide phosphates, which is consistent with the energetic patterns observed here.

Interestingly, the presence of Mg^2+^ significantly modulates the energetic contributions of several residues. For example, the magnitude of Lys525 stabilization is reduced in the IF-ATP-Sub-Mg^2+^ system compared to Mg^2+^-free conditions, suggesting that direct electrostatic interactions between Lys525 and the nucleotide phosphates are partially redistributed when Mg^2+^ coordinates the phosphate groups. This redistribution of interactions highlights the cooperative nature of nucleotide stabilization within the binding pocket.

Overall, the residue-level decomposition reveals that ATP stabilization in PCAT1 is dominated by contributions from the Walker A motif, particularly Lys525, while residues in the signature motif provide secondary stabilization and Walker B residues primarily contribute to catalytic organization rather than direct nucleotide binding. These findings provide a detailed molecular explanation for the nucleotide-binding energetics observed in the FEP calculations and demonstrate the utility of the per-residue free energy decomposition approach for dissecting the energetic architecture of ABC transporter nucleotide-binding sites.

## Conclusion

In this work, we used extensive microsecond-scale molecular dynamics simulations together with free energy perturbation calculations to investigate the structural and energetic determinants of nucleotide binding in the protease-containing ABC transporter PCAT1. By systematically comparing multiple biochemical conditions—including the presence or absence of substrate peptides and Mg^2+^ ions—we characterized how ligand coordination influences conformational stability, nucleotide retention, and binding energetics.

Structural analyses reveal that substrate and Mg^2+^ act cooperatively to stabilize the inward-facing (IF) conformation of PCAT1. Systems containing both substrate and Mg^2+^ exhibit reduced global RMSD fluctuations and lower residue-level flexibility compared with systems lacking one or both ligands. Consistent with these observations, nucleotide-binding stability analyses demonstrate that ADP readily dissociates from the nucleotide-binding pockets in the absence of Mg^2+^, whereas Mg^2+^ coordination strongly stabilizes nucleotide retention throughout the simulations.

Thermodynamic calculations using free energy perturbation further show that ATP binding is highly favorable in the IF conformation, particularly when Mg^2+^ is present. These results highlight the critical role of divalent cation coordination in organizing the nucleotide-binding pocket and promoting energetically favorable nucleotide binding.

To further dissect the molecular origins of nucleotide stabilization, we introduced a residue-level free energy decomposition approach that quantifies the energetic contribution of individual residues to nucleotide binding. This analysis reveals that the Walker A lysine (Lys525) provides the dominant stabilizing interaction with ATP across all conditions, while neighboring Walker A residues contribute additional stabilizing interactions with the phosphate groups. In contrast, acidic residues in the Walker B motif exhibit positive interaction energies, consistent with their established catalytic roles rather than direct nucleotide stabilization.

Together, these results provide a detailed molecular picture of how ligand coordination, catalytic-site residues, and global transporter conformations jointly determine nucleotide stability in PCAT1. More broadly, the residue-level free energy analysis introduced here offers a powerful framework for dissecting the energetic architecture of nucleotide recognition in ABC transporters. Future studies integrating simulation with emerging cryo-EM structures and experimental mutagenesis will further clarify how cargo recognition and nucleotide binding are coupled during the PCAT1 transport cycle.

## Acknowledgement

This research was supported by the National Institute of General Medical Sciences (NIH grant R35GM147423 awarded to M.M.), the National Science Foundation (NSF grant CHE 1945465 awarded to M.M.), and the Arkansas Biosciences Institute. Computational resources were provided by the Texas Advanced Computing Center (TACC) at the University of Texas at Austin (Frontera) through LRAC allocation CHE21003 to M.M. The work also used Stampede at TACC and Bridges-2 at the Pittsburgh Supercomputing Center through allocation MCB150129 from the Advanced Cyberinfrastructure Coordination Ecosystem: Services & Support (ACCESS) program. Additional computational support came from the Arkansas High-Performance Computing Center, funded by multiple NSF grants and the Arkansas Economic Development Commission. We also acknowledge Drs. Shadi Badiee, Dylan Ogden, and Adithya Polasa for their continuous support and useful discussions.

